# Expansion of frozen hybrids in the guppy ectoparasite, *Gyrodactylus turnbulli*

**DOI:** 10.1101/2020.04.02.021329

**Authors:** Mateusz Konczal, Karolina J. Przesmycka, Ryan S. Mohammed, Christoph Hahn, Jo Cable, Jacek Radwan

## Abstract

Hybridization is one of the major factors contributing to the emergence of highly successful parasites. Hybrid vigor can play an important role in this process, but subsequent rounds of recombination in the hybrid population may dilute its effects. Increased fitness of hybrids can, however, be frozen by asexual reproduction. Here, we identify invasion of a “frozen hybrid” genotype in natural populations of *Gyrodactylus turnbulli*, a facultatively sexual ectoparasitic flatworm that causes significant damage to its fish host. We re-sequenced genomes of these parasites infecting guppies from six Trinidad and Tobago populations, and found surprisingly high discrepancy in genome-wide nucleotide diversity between islands. The elevated heterozygosity on Tobago is maintained by predominantly clonal reproduction of hybrids formed from two diverged genomes. Hybridization has been followed by spread of the hybrids across the island, implying a selective advantage compared to native genotypes. Our results thus highlight that a single outcrossing event may be independently sufficient to cause pathogen expansion.

## Introduction

Hybridization refers to a successful mating between individuals from two populations, which are distinguishable on the basis of one or more heritable characters and have divergent genomes (Arnold, 2004). It can be a potent evolutionary force, creating opportunities for adaptive evolution by increasing genetic variation on which natural selection can act (Arnold, 1997, 2004; Oziolor et al., 2019; Rieseberg et al., 2003; Taylor & Larson, 2019). The ecological and evolutionary consequences are potentially large and may include significant effects on the evolutionary trajectories of pathogenic organisms (‘parasites’ henceforth) (King, Stelkens, Webster, Smith, & Brockhurst, 2015).

This role is likely to increase in importance because human activities increase opportunities for hybridization by inducing environmental changes and incidental translocations (Brooks & Hoberg, 2007; Cable et al., 2017; Patz, Graczyk, Geller, & Vittor, 2000). Understanding the role of hybridization in parasite evolution is therefore essential if we are to predict hot spots of disease risk and emergence (Laine, Barrès, Numminen, & Siren, 2019).

Parasite strains of hybrid origin can gain an advantage because recombinants – individuals with shuffled genomes - produced after hybridization may show enhanced phenotypic characteristics compared to the parental generation, such as higher infectivity (Grigg, Bonnefoy, Hehl, Suzuki, & Boothroyd, 2001), expanded host range (Detwiler & Criscione, 2010), increased transmission potential (Volf et al., 2007) or increased tolerance to harsh environmental conditions (Laine et al., 2019). Hybridization, however, is often associated with asexual reproduction, which may in the long term preclude the initial benefits resulting from recombination between divergent genomes, since asexual lineages tend to be less adaptable to environmental change compared to sexually reproducing lineages (Neaves & Baumann, 2011). Asexual reproduction is thought to be enforced by Bateson-Dobzhansky-Muller incompatibilities accumulated in parental lineages (Janko et al., 2018; Neaves & Baumann, 2011). Asexual lineages frequently originate from hybridization events, for example in cases when the genomes of parental species are divergent enough to disrupt meiosis in hybrids, yet not diverged enough to seriously affect hybrid viability (Janko et al., 2018; Lamelza et al., 2019; Moritz et al., 1989). Alternatively, the hybrid origin of asexual animals can arise from hampered pairings of homologous chromosomes due to decreasing sequence homology (De Storme & Mason, 2014) or from asynchronous expression of genes brought together by hybridization (Carman, 1997).

Conversely, asexual reproduction may be selectively beneficial if clonal reproduction “freezes” heterozygosity that confers short-term evolutionary advantage (Neiman, Sharbel, & Schwander, 2014). Hybrids often show greater vigor, productivity, disease resistance, or overall increased Darwinian fitness (Kingsbury, 2009). Two alternative hypotheses may explain the existence of hybrid vigor (heterosis) (Lippman & Zamir, 2007). The overdominance hypothesis, attributes heterosis to superior fitness of heterozygous genotypes over homozygous wild types. The dominance hypothesis assumes that recessive deleterious mutations are to some extent masked in a heterozygous stage. Recent theoretical work indicates that dominance-induced heterosis might play an important role in evolution, especially if deleterious alleles are not effectively removed from populations as it is expected when populations are small (Gilbert, Pouyet, Excoffier, & Peischl, 2020; Kim, Huber, & Lohmueller, 2018; Zhao & Charlesworth, 2016). This mechanism may be particularly relevant to parasite evolution. Many parasite populations are small or experience frequent bottlenecks, therefore are prone to the accumulation of deleterious mutations (M. Lynch, Butcher, Bürger, & Gabriel, 1993), but also to elevated homozygosity due to inbreeding. In such a case, one could expect higher fitness of hybrid parasites. Successful parasite strains originating from a hybridization event have been documented in several species, including *Batrachochytrium dendrobatidis* (Farrer et al., 2011; Greenspan et al., 2018), *Toxoplasma gondii* and *Sarcocystis neurona* (Grigg et al., 2001; Wendte et al., 2010). However, because in all these cases some degree of recombination occurred, the relative role of heterozygosity versus recombination between highly divergent genomes remains unresolved.

Here, we report frozen genome-wide heterozygosity in wild *Gyrodactylus turnbulli*, a common guppy parasite and well-known model species in epidemiological and eco-evolutionary research (e.g. Houde & Torio, 1992; Jacquin et al., 2016; Phillips et al., 2018; Reynolds, Arapi, & Cable, 2018; van Oosterhout, Harris, & Cable, 2003). Previous laboratory work demonstrated that *G. turnbulli* from mixed strain infections often hybridize and have higher fitness (more parasites per host, longer infection, but not increased host mortality), suggesting that in fact hybrid individuals might have increased fitness, possibly due to heterosis or recombination (Schelkle, Faria, Johnson, van Oosterhout, & Cable, 2012). However, the effect of hybridization on population structure and dynamics of infections in wild populations has not been investigated to date (Xavier et al., 2015). We analyze thirty whole genomes of *G. turnbulli* originated from six natural populations from Trinidad and Tobago. On Tobago we identified a widespread, recently formed “frozen hybrid” genotype, suggesting the potential importance of heterosis in clonal invasions of *G. turnbulli* and highlight that a single hybridization event can lead to the emergence of highly successful parasites, despite suppressing recombination.

## Materials and Methods

### Field collections

During two expeditions (2017 and 2018), guppies (*Poecilia reticulata*) were sampled from six locations on Trinidad and Tobago (Trinidad: Aripo River [ARI], Caura River [CAU], Lopinot River [LOP]; Tobago: Dog River [DR], Health Center [HC], Spring Site [SPS]). Fish were transported to our field station in Charlotteville, northeast Tobago, where each population was kept in a separate aquarium. Over four weeks, guppies were subsequently screened for ectoparasites under a dissecting microscope. If gyrodactylids were observed, a single worm was allowed to move to a naïve, anaesthetized fish, obtained from *Gyrodactylus-free* mesocosms established by Phillips et al. (2018). The transfer was closely monitored under the microscope to ensure movement of just a single worm. Fish experimentally infected in this way were kept in separate tanks in order to establish strains from which we could obtain enough DNA required to prepare libraries for sequencing. Fish were screened every 2-3 days to verify the presence of gyrodactylids, and hosts were euthanized with an overdose of MS-222 8-13 days post-infection. The number of worms was counted, samples were preserved in 97% molecular grade ethanol and transported to Poland. Life expectancy for *G. turnbulli* was estimated to 4.2 days (Scott, 1982). The first offspring is born approx. 1 day after the birth of the parent, and subsequent offspring are born at 2-2.5 days intervals (Scott, 1982). We thus expect that most individuals develop from the first or second births. The first-born always develops clonally, while the second-born daughter originates from parthenogenesis (Bakke, Cable, & Harris, 2007; Cable & Harris, 2002). Even though it is unclear which mechanism restores the diploid component during parthenogenesis (Cable & Harris, 2002), it has been suggested that heterozygosity is maintained during that process, as reported for *G. salaris*. Third- and subsequent born daughters may result from outcrossing, selfing, or parthenogenesis. Given our experimental design we expect that the majority of gyrodactylid individuals on each experimentally infected fish developed from first or second births and should have preserved founder’s genomes. Samples with at least 10 worms were used for DNA extraction and for species identification as described below. Parasites collected in 2017 and identified as *G. bullatarudis* were reported in a separate study (Konczal et al., 2020), while this work focused solely on *G. turnbulli*.

### DNA extraction, libraries preparation and sequencing

DNA was extracted from pools of individuals (‘strain’ henceforth) derived from a single worm collected from a single experimentally infected fish using MagJET Genomic DNA kits (Thermo Scientific). DNA concentration was measured using Qubit High Sensitivity reagents. *Gyrodactylus* were identified to species using Sanger sequencing of a 262 bp fragment of the mitochondrial protein coding COII gene (Xavier et al., 2015). Sequences were aligned together with records downloaded from the NCBI Genbank, and a Neighbor Joining tree was reconstructed with MEGA X (Kumar, Stecher, Li, Knyaz, & Tamura, 2018) to determine species. Thirty strains were selected for library preparation (Nextera DNA Flex Library Prep Kit, Illumina) based on DNA quantity and population of origin. Subsets of strains were sequenced in 2018 with HiSeq2500 (2 x 100 bp). All other strains were sequenced in 2019 on the NovaSeq6000 platform in 2 x 150 bp mode (see Supplementary Table 1 for more details).

### SNP calling and population genomic analyses

The quality of raw reads was assessed with Fastqc (Andrews & Babraham Bioinformatics, 2010). Reads were then filtered with trimmomatic (version 0.36; Bolger, Lohse, & Usadel, 2014) and mapped using bwa mem (version 0.7.10-r789) to the reference nuclear genome (Hahn et al., in prep.), which was assembled from individuals from the *Gt3* laboratory strain kept at Cardiff University. Mapping results was visualized with Qualimap (García-Alcalde et al., 2012) and duplicated reads were marked with picardtools (version 2.18.5-6). SNPs were then called with samtools mpileup (version 1.6.0, options -R -C50 -t DP, ADF, ADR). Based on inspection of the distributions of diagnostic statistics, we removed SNPs with DP > 1200 or DP < 400 (for all samples together), with QUAL < 30, with MQ < 30 or in 5 bp distance from identified indels.

We then used SNPs to calculate PCA with plink (version 1.90, Purcell et al., 2007) using only SNPs with minor allele frequency greater than 0.05. The same dataset with additional sample representing reference genotype (all genotypes encoded as reference homozygous) was used to calculate distance matrix (in one minus identity-by-state units) with plink and to construct a Neighbor Joining tree using the ape package in R. In addition, we ran PCA analyses with SNPs, that were previously pruned for Linkage Disequilibrium. We first pruned SNPs with-indep-pairwise 100 10 0.1 option in plink, and then used the same program to calculate PCA (D. H. Alexander, Novembre, & Lange, 2009).

Vcftools (version v0.1.12b; Danecek et al., 2011) was used to calculate nucleotide diversity (π) and genetic divergence between populations (F_ST_) in 25 kb non-overlapping windows, as well as inbreeding coefficient (F_IS_, option–het) per strain. To calculate F_IS_ per strain we considered only SNPs polymorphic within population of origin. The fraction of heterozygous sites per genomic window was calculated using a custom script and our estimates of genome wide heterozygosity per strain were inferred as the average of these fractions across all genomic windows. For each strain we also calculated divergence form the reference genome in 25 kb non-overlapping windows, by counting the number of non-reference variants (alternative homozygous sites were counted as one and reference/alternative allele heterozygous sites were counted as half) and dividing it by a window length. Windows shorter than 12.5 kb (ends of scaffolds and scaffolds shorter than that length) were excluded from all analyses.

### Mitochondrial genome assembly and mtDNA analyses

To reconstruct phylogenetic relationships between strains we reconstructed the entire mitochondrial haplotype for strain G126 using MITObim (*de novo* approach of Hahn, Bachmann, & Chevreux, 2013) with COII fragment downloaded from NCBI (accession KP168411.1) as an initial bait. We then mapped reads from all sequenced strains to that sequence using bwa mem with default parameters. Results were inspected manually using IGV. Duplicates were marked with picardtools and SNPs were called using mpileup and bcftools assuming haploid samples. SNPs with quality lower than phred 30 were discarded from analyses. The nucleotide sequence for each strain was reconstructed with bcftools consensus. Sequences were then aligned and used for phylogenetic analyses.

The evolutionary relationship between strains was inferred using ML with a GTR model, implemented in MEGA X Software (Kumar et al., 2018). The same software and model were used to calculate mean evolutionary distances between groups. BEAST2 (v2.6.1; Bouckaert et al., 2014) was then used to perform Bayesian phylogenetic analysis of mitochondrial genomes and to estimate time to most recent common ancestor (TMRCA). TMRCAs of nodes of interest were estimated based on 10,000,000 generations, logging every 1,000 steps, with the first 20% generations discarded as a burn-in. We used a Yule Model as prior, HKY substitution model, and strict clock model. Divergence time estimates were based on 13.1% divergence per million years estimated for *G. salaris* (see Kuusela et al., 2007).

## Results

A total of thirty *G. turnbulli* strains from three Trinidadian (Aripo, Caura and Lopinot Rivers) and three Tobagonian populations (Dog River, Health Center and Spring Site; Figure 1A, details in Supplementary Table 1) collected in two consecutive years were sequenced to an average coverage of 38.7 per strain. Sequencing reads were then mapped to the reference nuclear genome and used for SNP calling. We identified 776,881 single nucleotide- and 161,255 indel-polymorphisms. Principal Component Analyses of SNP genotypes clearly separated Trinidad from Tobago, while differentiation between local populations was present only in some cases (Figure 1B). In particular, strains from Health Center and Dog River (Tobago) showed virtually no differentiation and, in contrast to the Trinidadian populations, do not form monophyletic groups according to the populations of origin (Figure 1C). The Trinidadian strains were more similar to the reference genome, obtained from a strain on commercial guppies, *Gt3*, than strains collected from Tobago (Figure 1C). To remove the potential effect of linkage on population differentiation, we pruned linked SNPs, but the general pattern remained the same (Supplementary Figure 1). We calculated F_ST_ between Trinidad and Tobago populations in 25 kb non-overlapping windows and found that 90% of the windows show F_ST_ values between 0.25 and 0.55 (mean F_ST_ = 0.42; Supplementary Figure 2). Pairwise F_ST_ values were higher among Trinidadian populations (Supplementary Figures 3) than among populations from Tobago (Supplementary Figures 4). To resolve whether moderate F_ST_ between islands resulted from moderate diversification between or from elevated levels of genetic variation within populations, we calculated nucleotide diversity across Trinidad and Tobago strains separately. Nucleotide diversity on Tobago was 2.4 times greater than that on Trinidad (0.0028 and 0.0012 respectively, Figure 1D). The difference does not result from high differentiation among Tobagonian populations: nucleotide diversity was also higher within Tobagonian populations, compared with those from Trinidad (Supplementary Figure 5).

**Figure 1.**
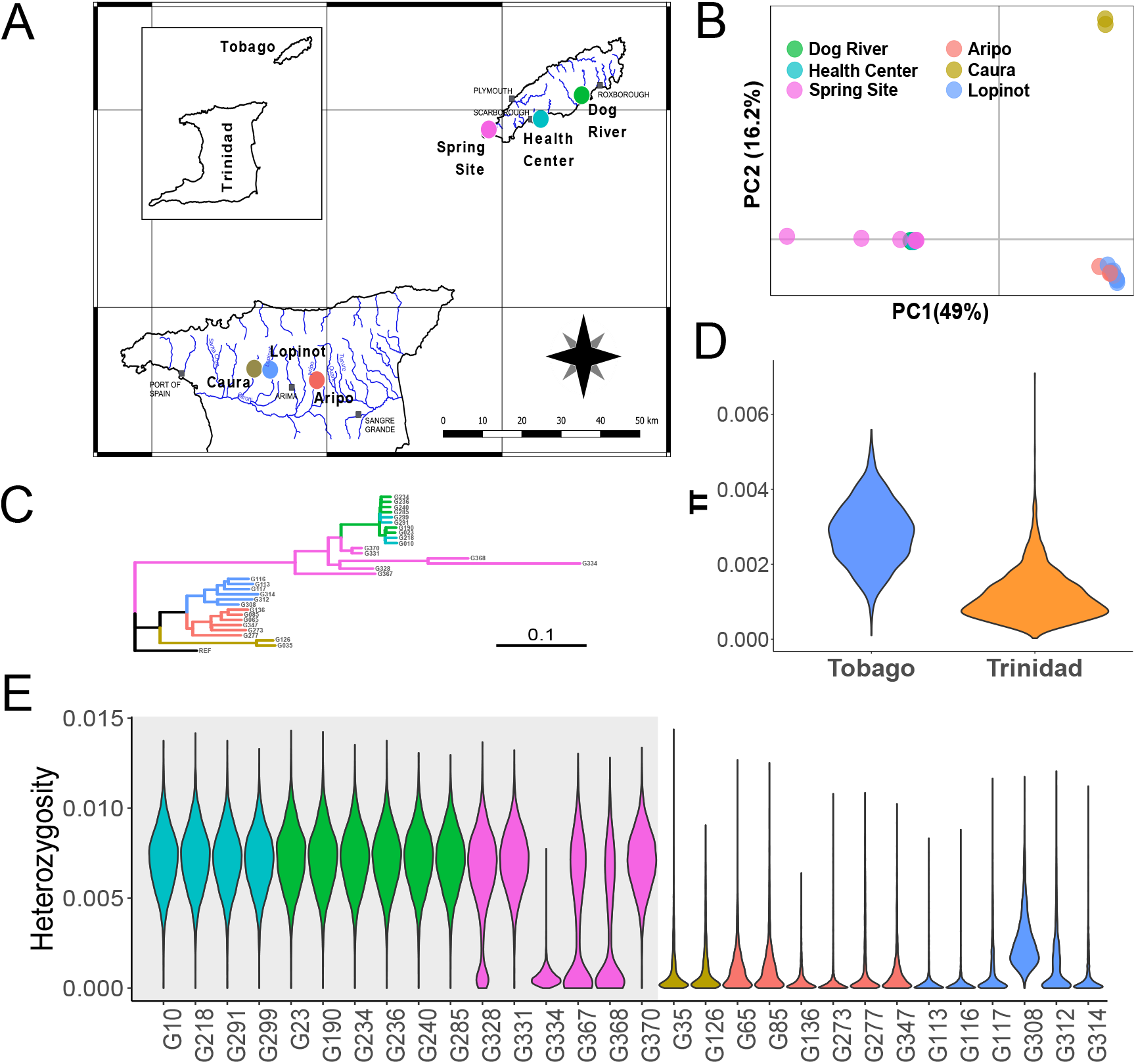
Patterns of genetic variation in *Gyrodactylus turnbulli* populations from Trinidad and Tobago. **A:** Map of Trinidad and Tobago with sampling sites. **B:** Principal Component Analyses of *G. turnbulli* strains. PCA based on 673,104 SNPs which passed filtering criteria. SNPs were identified within 30 isolates from three Trinidadian and three Tobagonian populations. **C:** Neighbor Joining tree (Saitou and Nei 1987) constructed using distance matrix, with one minus identity-by-state units, calculated based on 673,104 SNPs. Populations are represented by different colours. The reference sequence (obtained from a strain on commercial guppies) is shown in black. **D:** Distribution of nucleotide diversity (π) calculated across the genome in 25 kb non-overlapping windows. **E:** Distribution of heterozygosity calculated in 25 kb non-overlapping windows for each strain collected from Trinidad (white background) and Tobago (shaded background) populations.

Increased nucleotide diversity on Tobago was mainly driven by high heterozygosity of most individuals (Figures 1D-E). Similarly, inbreeding coefficient, F_IS_, was negative for most of the strains from Tobago, suggesting over-representation of heterozygous positions (Supplementary Figure 6). In contrast, most of the Trinidadian strains showed positive values of F_IS_.

High heterozygosity was observed in all strains from the Tobago populations Health Center and Dog River. The third Tobago population (Spring Site) showed a mixture of patterns. One Spring Site strain (G334) had low level of heterozygosity, reminiscent of the general pattern observed in Trinidadian strains. Three other strains (G328, G367 and G368) showed bimodal distribution, with genomic regions presenting either high or low level of heterozygosity. Finally, two Spring Site strains (G331 and G370) contained only highly heterozygous regions, similar to the pattern observed in Health Center and Dog River (Figure 1E).

The majority of positions found polymorphic among Tobagonian strains (325,473 out of 498,111) were heterozygous in all Health Center and Dog River strains, as well as and two (G331 and G370) Spring Site strains. A total of 88% of these SNPs combined one allele from the reference genome and the alternative allele found in the Tobagonian G334 strain (Figure 2). Genome-wide pattern of divergence between re-sequenced strains and the reference genome demonstrated that strain G334 was the most divergent. Strains from Trinidad were consistently more closely related to the reference genome (Figure 1C, Supplementary Figure 7). Based on these patterns we infer that highly heterozygous Tobagonian strains consist of hybrids between two divergent haplotypes: one similar to the reference sequence and the second similar to the G334 sequence. The same pattern with shared heterozygous sites and high, genome-wide heterozygosity was observed in the Health Center and Dog River strains from 2017 and 2018, indicating that the hybrid population is stable. This implies an absence of recombination, as recombination would have fixed one of the two haplotypes in at least some genomic regions. Auxiliary analysis comparing heterozygosity across all contigs to that of a simulated, maximally heterozygous hybrid genome, created *in silico* from the reference sequence and the G334 strain, confirmed that no such homozygous regions are observed in these populations (Supplementary Figure 8). Furthermore, we compared genome wide nucleotide diversity of Tobagonian strains collected in 2017 and 2018 and did not find evidence for a substantial difference between time points (π_2017_ = 0.0018 vs π_2018_ = 0.0017). We also plotted relationships between heterozygosity calculated in non-overlapping windows for each pair of strains and did not find any indication for a reduction in heterozygosity in any strain (Supplementary Figure 9), suggesting no ongoing recombination.

**Figure 2.**
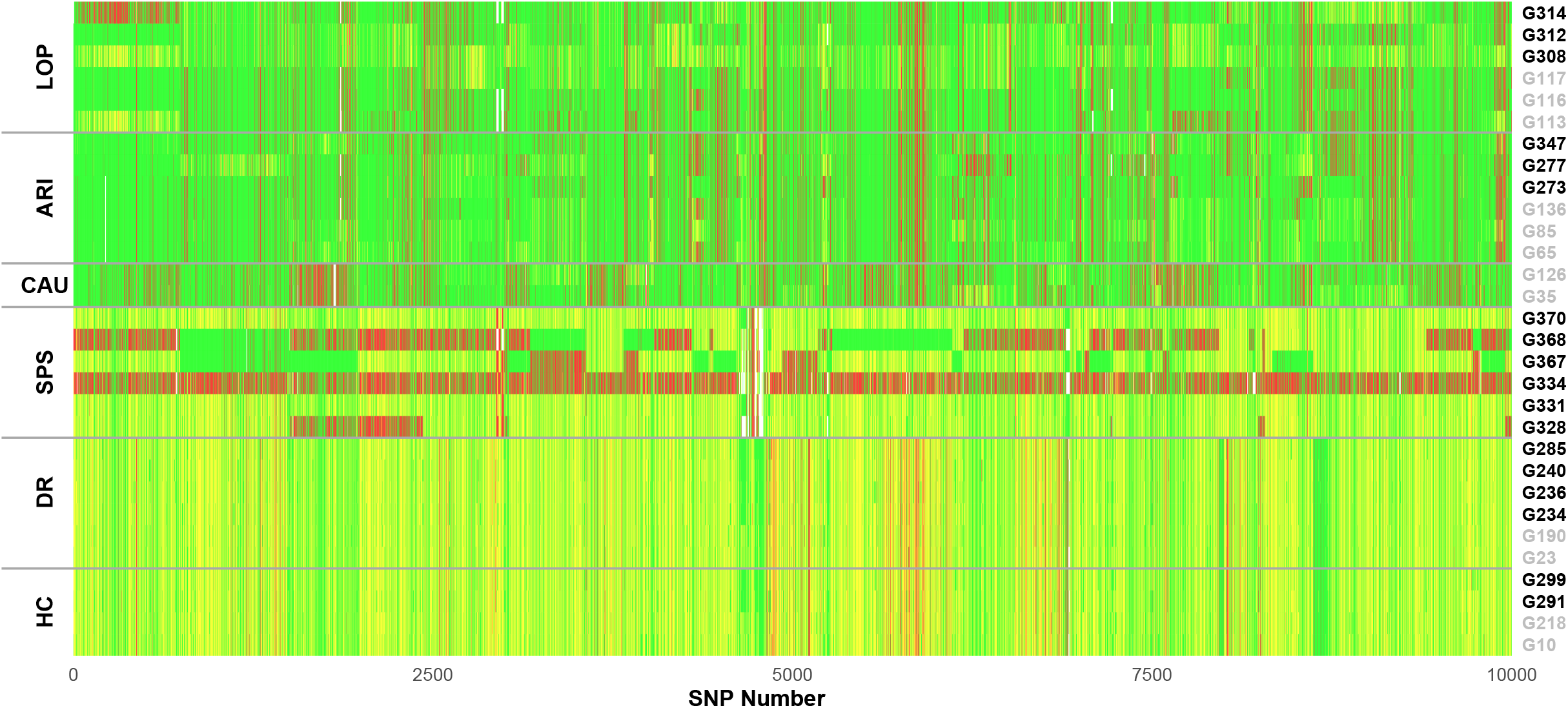
Graphical representation of genotypes in *Gyrodactylus turnbulli* strains. Genotypes identified as reference homozygotes are shown in green, alternative homozygotes in red, heterozygous in yellow. Strains collected in 2017 and 2018 are labeled in grey and black respectively, and those collected from different local populations are separated by horizontal grey lines. A subset of polymorphic sites (10,000 SNPs) from the longest scaffolds is shown.

In contrast, divergent haplotypes have clearly recombined in the Spring Site population. Three strains (G328, G367 and G368) show a mosaic pattern of the two haplotypes fixed in different genomic locations, with other parts of the genome still remaining heterozygous (Figure 2, Supplementary Figures 8 and 10). Signatures of sexual reproduction are also observed at least in some of the strains from Trinidad. For example, SNPs localized in the scaffold scf7180000443747 (first sequence in the reference genome) show admixture of another haplotype in strain G314 coming from Lopinot, and another two strains in Lopinot are heterozygous in this region (Figure 2). Another strain from the same population (G308) has negative F_IS_ (Supplementary Figure 6) and slightly elevated genome-wide heterozygosity (Figure 1D), although the effect is weaker compared to strains from Tobago. Nevertheless, heterozygous sites are on average rare in Trinidadian strains.

Clustering analyses (Figures 1B, 1C) and F_ST_ results (Supplementary Figure 4) based on nuclear SNPs suggest low divergence between samples identified as hybrids. We explored it further by calculating divergence in the haploid mitochondrial genomes. One Trinidadian strain (G126) was used for mitochondrial genome assembly, and then reads from all other strains were mapped to that genome for haplotype reconstruction. The resulting mt genome spans 14,159 bp. We found very little divergence between strains identified as hybrids, compared to divergence amongst other samples (Figure 3, Supplementary Figure 11). Putative hybrid strains from Health Center, Dog River and Spring Site populations were found more closely related to Trinidadian strains than to remaining SPS strains, confirming introgression inferred from genomic data. Within hybrids (Health Center, Dog River two hybrid strains from Spring Site) the mean divergence between populations was 0.02%. In contrast, the three other strains collected from Spring Site had strongly divergent mitochondria - the nucleotide divergence between the two Spring Site groups was 1.58%. To independently verify low divergence between Tobagonian hybrids and strains from Trinidad we sequenced a 262 bp fragment of COII from 81 samples collected by us on Trinidad and Tobago. The results confirmed little divergence between all Tobagonian samples except for some derived from the Spring Site population (Supplementary Figure 12).

**Figure 3.**
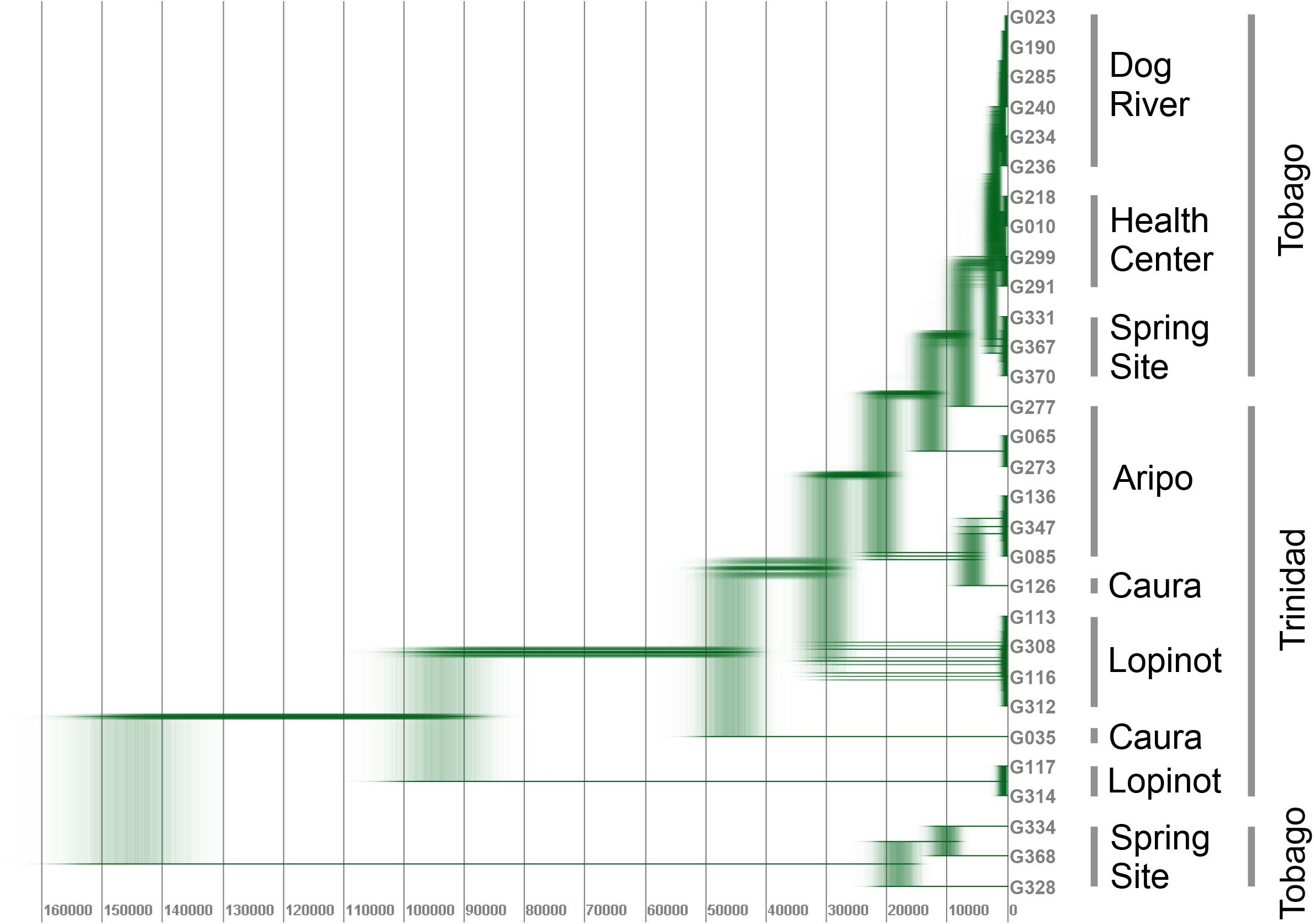
Phylogenetic relationship between *Gyrodactylus turnbulli* strains, reconstructed based on mitochondrial genome sequences (14,159 bp). The trees were derived via a Bayesian phylogenetic method, using 10,000,000 generations of MCMC samples, with HKY substitution model and strict clock model, assuming 13.1% divergence per million years. The estimated time before present (in years) for tree nodes is shown at the bottom of the tree.

Lack of recombination in Tobago hampers identification of fast evolving regions in the genome, because of strong linkage between sites. However, there was no evidence that the Trinidadian populations experienced clonal expansion, and therefore rapidly evolving genes can be potentially identified. For this reason, we tested for correlations between nucleotide diversity calculated in windows separately for Tobagonian hybrid populations and for all Trinidadian strains. We used nucleotide diversity calculated jointly for Health Center and Dog River strains as a proxy for the long-term evolution of genomic regions (divergence between two haplotypes forming hybrids). Genomic regions in which the nucleotide diversity calculated jointly for all strains from Trinidad deviates from predictions based on this long-term evolutionary rate are therefore candidates for recent rapid evolutionary change between years or populations. The divergence calculated in 25 kb windows were highly correlated between Tobagonian hybrid populations and Trinidadian populations, while few windows demonstrated elevated (according to overall trend) diversity in Trinidad (Figure 4A). One scaffold (scf7180000445747; 19.4 kb long) showed extensive divergence between Trinidadian strains collected in 2017 and 2018 (Figure 4B). It contains 7 protein coding genes, including putative methylotransferase NSUN5, Endonuclease/exonuclease/phosphatase family domain-containing protein 1, synaptogyrin and prohibitin.

**Figure 4.**
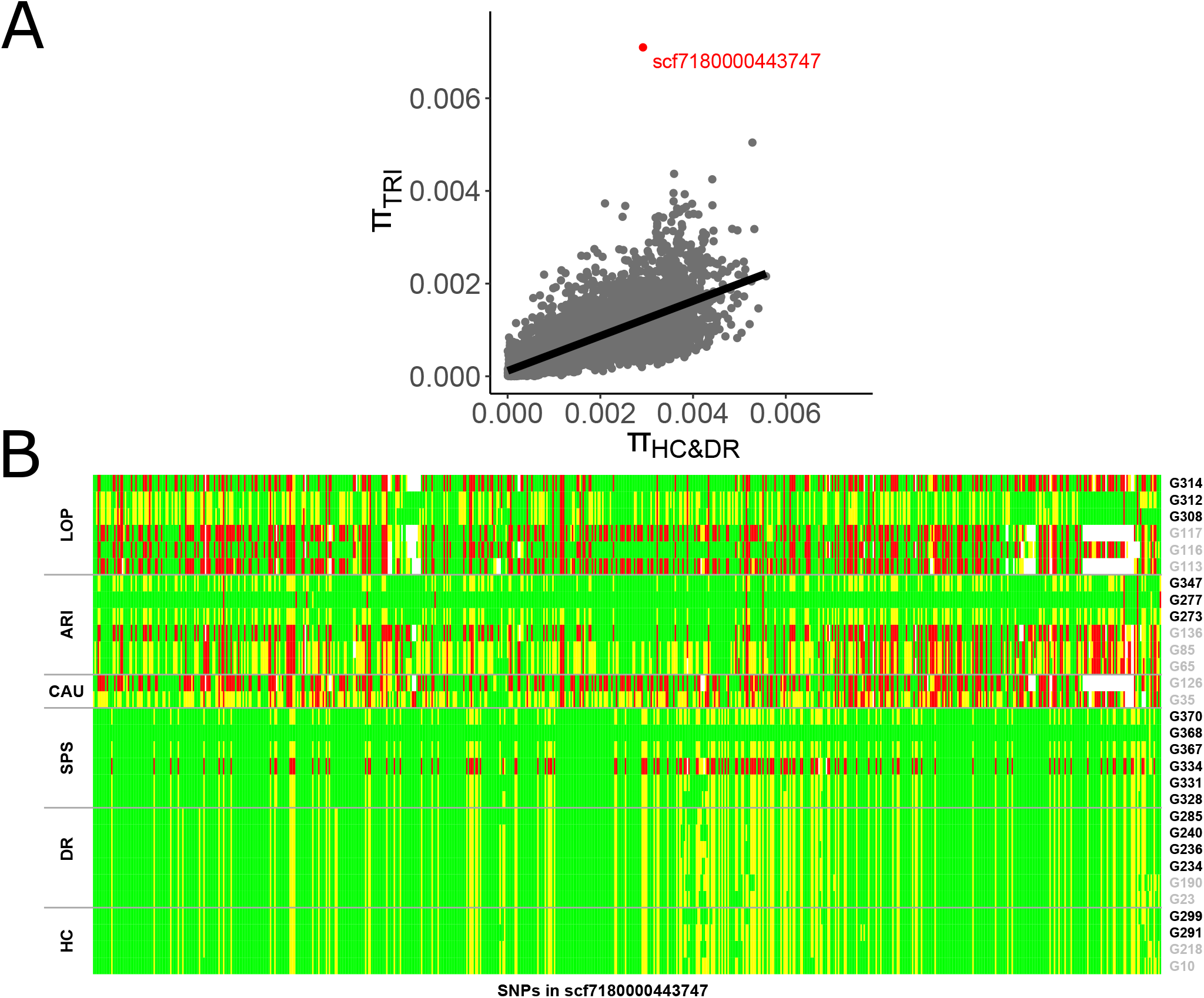
Genetic variation in genomic region with elevated nucleotide diversity in populations from Trinidad. **A:** Correlation between nucleotide diversity calculated jointly for Health Center and Dog River populations and nucleotide diversity calculated jointly for all strains from Trinidad. Black line represents linear regression (y = 0.00 + 0.38*x). Scaffold scf7180000443747 is shown in red. **B:** Graphical representation of genotypes in polymorphic sites of *Gyrodactylus turnbulli* localized in scaffold scf7180000443747 (707 SNPs). Genotypes identified as reference homozygotes are shown in green, alternative homozygotes in red, heterozygous in yellow. Strains collected in 2017 and 2018 are shown in grey and black respectively, and those collected from different local populations are separated by horizontal grey lines.

## Discussion

We examined whole genomes of thirty *G. turnbulli* strains, sampled from two Caribbean islands in two consecutive years, providing insight into the evolution of their genomic diversity. On Tobago, parasite populations tend to have much higher nucleotide diversity than those on Trinidad. This higher diversity was caused by the presence of highly heterozygous strains in all studied Tobagonian populations (50-100% of individuals per population). These strains shared most of the heterozygous loci, with a repeatable pattern of having one reference-like allele and the other allele present in a homozygous state only in one strain from Tobago (G334). This pattern led us to conclude that these highly heterozygous strains comprise hybrids of two divergent lineages, one of which is close to a strain isolated from ornamental guppies (Trinidad-like, used for the reference genome assembly), and the second which is close to the Tobagonian G334 strain. In the Spring Site population (Tobago), we observed two distinct mitochondrial lineages (diverged 145,000 years ago based on 13.1% divergence per Mya, (Kuusela et al., 2007), 95% HPD: 133,900 – 155,700). Most likely they represent mitochondrial genomes originating from two parental populations that formed Tobagonian hybrids. The mitochondrial divergence between these two lineages was much larger than between Tobagonian hybrids and Trinidadian populations (7,600 years, 95% HPD: 5,200 – 9,800 years), demonstrating that hybrids inherited mitochondria from Trinidad-like ancestors. Initial hybridization must have been followed by reproduction without recombination, otherwise recombination would have produced both homozygous and heterozygous genotypes at least in some of the genomic regions (as observed in two strains at Spring Site). In contrast, in most Tobagonian strains heterozygosity was elevated genome-wide (Figure 1D), heterozygous genotypes were very similar for all samples (Figure 2), suggesting that these genomes represent “frozen hybrids”. Indeed, our analyses imply that no recombination occurred since the hybrids were formed. If recombination happened before expansion of the hybrid genotype, some regions with loss of heterozygosity would be found and shared for all HC and DR strains. We did not find any evidence for loss of heterozygosity in HC nor DR (Supplementary Figure 8). In contrast to HC and DR, patterns suggesting recombination were obvious in some strains from the Spring Site population. Strains G328, G367 and G368 showed non-identical patterns of heterozygosity across the nuclear genome, consistent with recent independent outcrossing/selfing events followed by recombination. Two of these three Spring Site strains (G328, G368) have a mitochondrial genome significantly divergent from other hybrids, suggesting that subsequent rounds of outcrossing occurred between hybrid and non-hybrid individuals, the latter carrying the divergent mtDNA. On Trinidad, a single individual showed slightly elevated heterozygosity, possibly as a result of sexual reproduction between two moderately diverged strains (G308, Figure 1D, Supplementary Figure 6), and indeed another individual from the same population showed a pattern consistent with being a subsequent generation recombinant (G312). Thus, in contrast to a set of pure hybrid populations we found on Tobago (Dog River and Health Centre), recombination is not completely absent in other populations both on Tobago (Spring Site) and on Trinidad. This evidence is consistent with the data from laboratory cross infections with *G. turnbulli*, which demonstrated that 4-11% of individuals in mixed infections are hybrids with ancestors from different strains used in the experiment (Schelkle et al., 2012).

Not only sexual reproduction, but also most forms of parthenogenesis involve meiotic recombination, which is expected to lead to loss of heterozygosity spanning large genomic regions. However, automictic parthenogenesis (automixis) with central fusion can freeze heterozygosity in spite of recombination.(Engelstädter, 2017; Jaron, Bast, Ranallo-Benavidez, Robinson-Rechavi, & Schwander, 2018) Under automixis with central fusion the probability of recombination increases with distance from the centromere, and particularly regions distal to the centromere are affected if recombination does occur. While, we are not able identify centromeres/telomeres in the reference genome of *G. turnbulli*, we did not find any evidence for large genomic regions with loss of heterozygosity (Supplementary Figures 11 and 12). Thus, either Tobagonian hybrids (i) reproduce only mitotically, (ii) recombination is strongly suppressed during automictic parthenogenesis, or (iii) recombinants are removed by selection leading to clonal-like evolution. Our data suggest that the inferred, clonally-like reproducing hybrids spread relatively recently across Tobago, as indicated by high similarity of their mitochondrial genomes, corresponding to 2,700 (95% HPD: 1,500 – 4,000) years of divergence (Figure 3). The divergence time is based on the indirectly derived substitution rate in a single mitochondrial locus of *Gyrodactylus salaris* (13.1% per Mya; Kuusela et al., 2007). Other indirect estimates suggest lower (5.1% per Mya; Hahn et al., 2015) or higher substitution rates (up to 20.3% per Mya; Meinilä et al., 2004) in *Gyrodactylus* species, and therefore divergence dating should be taken with caution. Nevertheless, low nucleotide divergence between mitochondrial genomes indicates a recent origin of the hybrids, and their widespread occurrence across Tobago provides evidence for their evolutionary success. Hybrid genomes that were most widespread on Tobago did not show any signs of recombination, and apparently remained stable, as would be expected under predominantly clonal evolution.

What evolutionary forces are responsible for the apparent success of clonal hybrids on Tobago? Tibayrenc and Ayala (2012, 2017) proposed that a predominance of clonal evolution is a common adaptation to parasitism in micropathogens. Their hypothesis considers the total set of reproductive strategies used by pathogens to escape ‘recombination load’. In contrast, the ‘epidemic clonality model’ (Smith, Smith, O’Rourke, & Spratt, 1993) advocates for the occurrence of occasional bouts of clonally reproduced strains in otherwise recombining species. Ephemeral clones are frequently replaced by other genotype combinations, maintaining a dominant role of sexual reproduction in evolution. Complex patterns of outcrossing events in *G. turnbulli* populations seem to be consistent with an ‘epidemic clonality model’ of evolution.

In the case of *G. turnbulli* a bout of clonality is associated with hybridization between two, relatively divergent strains, confirming that hybridization of parasites can lead to the emergence of highly successful, and potentially virulent strains. The potential to increase the virulence of *G. turnbulli* by combining different genomes was suggested in an earlier report of laboratory experiments, in which infection with two inbred stains led to higher parasite load compared to single strain infections (Schelkle et al., 2012). However, the genomic divergence between these strains was unknown and, as discussed by the authors, this result could be due to competition between clones leading to the evolution of increased virulence (Schelkle et al., 2012).

Human activity, migrations and climate change shift the geographic distribution of many species (Brooks & Hoberg, 2007; Cable et al., 2017; Lafferty, 2009), which increases the probability of secondary contact between former distinct populations, potentially leading to hybridization followed by adaptive evolution. Parasites are often transported with their hosts, and even if hosts do not contribute to genetic pools of local populations, hitchhiking parasites can do so. This could be the case for *G. turnbulli*. Guppies are commonly kept in aquaria, and *G. turnbulli* are regularly reported in aquarium populations (Maceda-Veiga & Cable, 2019) from where they can potentially be released to nature. Alternatively, hybrids might have appeared naturally. Our approximate dating of divergence of mtDNA between hybrids from Trinidad and Tobago is consistent with the end of the last glacial maximum, when a land bridge that probably existed between the islands during glaciation-related sea level decline was disrupted again (H. J. Alexander, Taylor, Sze-Tsun Wu, & Breden, 2006; Lambeck, 2004). This might have facilitated migration of guppies between the islands, and possibly other fish species capable of spreading *G. turnbulli* (Cable et al., 2013; T. A. King & Cable, 2007).

Our recent study on the natural populations of another monogenean parasite infecting guppies (*Gyrodactylus bullatarudis*) suggested that recent hybridization has also played a predominant role in shaping genetic variation, but in that case in populations from Trinidad (Konczal et al., 2020). Hybridization between two diverged lineages and its subsequent recombination resulted in a mosaic genome composition of *G. bullatarudis*. Around half of the genome originates from one or another lineage. High divergence between recombined lineages and relatively little divergence between local populations led us to conclude that recent hybridization was followed by rapid expansion of a recombinant *G. bulllatarudis* strain. Similar patterns of successful admixed hybrid genotypes have been reported for many other parasites, including schistosomes (Kincaid-Smith et al., 2019; Platt et al., 2019), *Trypanosoma* (see Tihon et al., 2017) and *Leishmania* (see Rogers et al., 2014) species, as well as Dutch elm disease pathogens (Brasier, 2001; Hessenauer et al., 2020) and other fungi pathogens (Mixão & Gabaldón, 2018) highlighting the role of recombination between divergent genomes in parasite evolution. Our analysis of *G. turnbulli* genomes indicates, that unlike its congener *G. bullatarudis*, natural expansion occurred without prior recombination. This scenario is similar to suggestions for *Gyrodactylus salaris* - a significant pathogen of Atlantic salmon. The Baltic-salmon specific *G. salaris* was proposed to have originated from a single hybridization event between two clades, followed by the clonal reproduction of ‘frozen hybrids’, however this suggestion was based on analyses of a single anonymous nuclear marker (Kuusela et al., 2007). Therefore, not only clonal expansion of hybrids, but also linkage to lethal variants for both alleles (balanced lethal system), or epistatic interactions could potentially explain the pattern observed in *G. salaris*. Our genome-wide data suggest that clonal expansion of ‘frozen hybrids’ can indeed take place amongst gyrodactylids.

Similar observations in other pathogens also suggest that ‘frozen hybrids’ might have a significant evolutionary advantage. An interesting example comes from a common fungal parasite of amphibians, *Batrachochytrium dendrobatidis*, in which asexually reproducing hybrids have higher virulence than both parental strains in some host species (Farrer et al., 2011; Greenspan et al., 2018). That hybrid is highly heterozygous and widespread. Most likely contact between previously genetically isolated allopatric populations of the fungus facilitated hybridization, resulting in generation, spread and invasion of the hypervirulent strain (Farrer et al., 2011). However, the strain is not completely clonal and diversification can proceed by either mitotic or sexual recombination. Similarly, in *Toxoplasma gondii* and *Sarcocystis neurona* outcrossing can precede a disease outbreak (Grigg et al., 2001; Wendte et al., 2010), but subsequent selfing events were responsible for epidemic expansions, and, as pointed out by the authors, it is not clear whether an out-cross is independently sufficient to cause an epidemic (Wendte et al., 2010). Our data are therefore unique, in demonstrating parasite expansion despite complete genome-wide suppression of recombination, implying that ‘frozen hybrids’ might gain advantage from masking recessive, deleterious mutations or from widening ecological niche.

Parasites may be particularly prone to accumulation of deleterious mutations in their genomes (Criscione & Blouin, 2005). In general, the rate of accumulation depends primarily on the effective population size. In small populations, random genetic drift can overpower selection making it easier for deleterious mutations, particularly those with small effects, to become fixed (Kimura, Maruyama, & Crow, 1963). This can cause mutational melt-down leading to population extinction (Michael Lynch & Gabriel, 1990). Several features of macroparasite life cycles can act in concert to reduce effective population sizes and increase the risk of such melt-down (Criscione & Blouin, 2005). In particular, frequent bottlenecks associated with colonization/recolonization of new hosts can significantly reduce the efficiency of selection in removing mildly deleterious mutations from populations. Additionally, observations of *G. turnbulli* in the wild suggest that census population sizes are often small and fluctuating over time (Stephenson, van Oosterhout, Mohammed, & Cable, 2015), which is consistent with the low heterozygosity and generally positive inbreeding coefficient in the non-hybrid populations that we investigated (Figure 1E, Supplementary Figure 6). If this leads to homozygosity for a number of recessive or partially recessive mutations, hybridization between divergent strains carrying those mutations at different genomic locations would result in hybrid vigor. Alternatively, the advantage of hybridization can be due to alleles carried by heterozygotes being adapted to different, and a wider range of, environments. In the case of parasites, divergent alleles may each allow infection of different host species or genomes, allowing heterozygotes to infect a wider range of hosts. For example, laboratory experiments on tapeworms in sticklebacks showed that two genetic lines were only able to infect a single respective host species, whereas their hybrids could infect both species (Henrich, Benesh, & Kalbe, 2013). In the case of *G. turnbulli*, however, hybridization occurred among two strains infecting the same host species.

Clonal expansion of hybrids is likely facilitated by facultative sexual reproduction of *Gyrodactylus* species. In such systems, switching between facultative and obligate clonal reproduction does not appear to be evolutionarily constrained and can easily evolve when favoured by selection (Neiman et al., 2014). However, though it has been proposed that hybridization is the main mechanism responsible for evolution of asexual reproduction, its causality and underlying mechanisms are poorly understood (Janko et al., 2018; Neiman et al., 2014; Schön, van Dijk, & Martens, 2009; Xu, Innes, Lynch, & Cristescu, 2013). Mortiz et al. (1989) suggested that reproducing asexually results from the accumulation of incompatibilities between hybridizing species that disrupts cellular regulation of sexual reproduction. In fact, the majority of species which recently evolved asexual reproduction are of hybrid origin, although it concerns primarily vertebrates and it is not clear whether it applies also to other taxa (Neaves & Baumann, 2011). In the case of *G. turnbulli*, however, recombination is not completely suppressed, because it was observed in one of the hybrid populations on Tobago (Spring Site), where it might have occurred between hybrid and non-hybrid individuals. The fact that recombinant individuals are rare on Tobago might result either from rarity of recombination events, or from effective removal of recombinants from the population by selection. The disadvantage of recombinants may stem from restoring homozygotes for recessive deleterious mutations. Discrimination between these two possibilities ideally requires direct estimates of recombination rate, possibly based on changes in genotype frequencies over time in both hybrid and non-hybrid populations (Becheler et al., 2017).

Recombination can, at least to some degree, enable independent evolution of different genomic regions, allowing identification of genomic regions with elevated genomic variation between the two sampling years, possibly driven by selection. We therefore searched for such regions in Trinidad populations and found that one genomic region showed an extreme pattern of genetic diversity, with several interesting genes previously associated with modulating lifespan, stress resistance (Methylotrasferase NSUN5; Schosserer et al., 2015) and host-parasite interactions (Prohibitin; Jain et al., 2010). Future work including more individuals and experimental validation is needed to confirm involvement of these genes in coevolutionary dynamics.

Concluding, our population genomic dataset of *G. turnbulli* allowed us to infer that hybridization followed by clonal reproduction contributed to evolutionary success of these parasites on Tobago. It is likely that the evolutionary success stems from heterotic effects, which are frozen in hybrids. Benefits of such freezing may only be temporary, as the lack of recombination may hinder the pathogen in keeping up in the evolutionary arms-race with the host’s immune system. That recombination seemed to be maintained in Trinidadian populations suggests that it may be favoured in the long run as well. It would be interesting to see if after initial success, recombination would again be favoured, as might have been the case with *G. bullatarudis* (see Konczal et al. 2020). Our results highlight that a single outcrossing event is independently sufficient to cause pathogen expansion, thus emphasizing that such processes can result in emergence of invasive pathogens, some of which may be of public health and conservation concern.

## Acknowledgements

We thank K. Phillips for his help in improving this manuscript, N. Cook, A. Szubert-Kruszyńska, A. Sadowska-Konczal and staff of the Environmental Research Institute Charlotteville (ERIC), Tobago, for support in the field; P. Turpin of Man O’War Bay Cottages for renting us the field station; Tobago House of Assembly (THA) for granting permission to conduct field surveys; K. Dudek and J. Raubic for help in molecular laboratory. All animal work was approved by Cardiff Universities Animal Ethics Committee and followed UK Home Office Licence (PPL 303424). This work was supported by Polish National Science Center Fuga Grant UMO-509-2016/20/S/NZ8/00208. Computations were performed at the Poznan Supercomputing and Networking Center.

## Data accessibility

The raw sequences are available as FASTQ files in the GeneBank (BioProject accession no. PRJNA603444).

## Author Contributions

M.K. and J.R. designed research; M.K., K.J.P. and R.S.M. collected samples; C.H. provided reference genome; M.K. analysed data with suggestions from J.R., J.C. and C.H.; MK drafted manuscript, J.R. and J.C. contributed to manuscript writing. All authors corrected manuscript and approved the final version.

## Notes

### Competing Interest Statement

The authors have declared no competing interest.

